# Identifying endogenous peptide receptors by combining structure and transmembrane topology prediction

**DOI:** 10.1101/2022.10.28.514036

**Authors:** Felix Teufel, Jan C. Refsgaard, Marina A. Kasimova, Christian T. Madsen, Carsten Stahlhut, Mads Grønborg, Ole Winther, Dennis Madsen

## Abstract

Many secreted endogenous peptides rely on signalling pathways to exert their function in the body. While peptides can be discovered through high throughput technologies, their cognate receptors typically cannot, hindering the understanding of their mode of action. We investigate the use of AlphaFold-Multimer for identifying the cognate receptors of secreted endogenous peptides in human receptor libraries without any prior knowledge about likely candidates. We find that AlphaFold’s predicted confidence metrics have strong performance for prioritizing true peptide-receptor interactions. By applying transmembrane topology prediction using DeepTMHMM, we further improve performance by detecting and filtering biologically implausible predicted interactions. In a library of 1112 human receptors, the method ranks true receptors in the top percentile on average for 11 benchmark peptide-receptor pairs.

## 1 Introduction

Endogenous bioactive peptides are ubiquitous in higher organisms and involved in many physiological processes, ranging from controlling metabolism [1] to neural signaling [2]. Many of these peptides are secreted and exert their function by binding to membrane-expressed receptor proteins, such as G-protein coupled receptors (GPCRs) and are thus appealing for drug development [3]. While an increasing number of potential peptides are discovered through technologies such as mass spectrometry peptidomics [4], small open reading frame sequencing [5] and bioinformatics approaches [6], the number of known peptide-receptor pairs remains stagnant, with currently 488 human peptide-receptor interactions reported in the GPCRdb [7]. This is due to the fact that experimental validation of peptide receptors requires tedious experimental screening of cell lines and binding assays, which still scale poorly in light of the large search space of potential receptors. Estimates reported at least 1200-1300 surface-expressed receptors in human [8, 9] even when excluding many isoforms and proteins without manual evidence, making exhaustive screening infeasible.

A previous computational approach for peptide-receptor pairing [6] has used prior knowledge, structural analyses and machine learning to identify potential peptide receptors within the complete receptome, thereby reducing the number of receptors that need to be screened. Using comprehensive experiments, 17 endogenous peptides could thereby be paired within a single study. While this result presents a substantial advancement, it still relies on experiments for pairing at its core.

AlphaFold [10, 11] has shown state-of-the-art performance for predicting protein-peptide interactions [12, 13]. However, research has so far focused on peptide docking to mostly globular proteins, and peptide binding prediction accuracy was evaluated using a moderately imbalanced ratio of positives and negatives (1:5). In this work, we investigate the application of AlphaFold for the prediction of binding of peptides to membrane-bound receptors. This problem is characterized by a severe class imbalance, as only a few receptors out of hundreds of candidates are expected to truly bind a peptide of interest.

Another aspect specific to receptor interactions is the receptor’s transmembrane topology, which can be predicted directly from its amino acid sequence [14]. While AlphaFold has strong performance for folding transmembrane proteins [15], it has no explicit knowledge about the interaction constraints imposed by the membrane topology, which are relevant especially for complex prediction where the localization of the binding partner is known. As predicted AlphaFold binding might violate these imposed constraints, we explore the use of DeepTMHMM [16] to detect spurious binding and improve prediction performance.

## 2 Method

We approach the question of identifying endogenous peptide receptors as a ranking problem: Given a list of candidate receptors, an ordered list of all receptors should be produced with true receptors ranked at the top. For a library of candidate receptors and a peptide of interest, we apply AlphaFold-Multimer [11] to predict a protein-peptide complex for each receptor. Additionally, we use DeepTMHMM [16] to predict the transmembrane topology of each receptor (Figure 1). DeepTMHMM is a protein language model [17] based predictor that only takes the amino acid sequence as input, thereby providing us with information orthogonal to the structure predicted by AlphaFold. For each residue, it predicts an {Intracellular, Extracellular, Transmembrane, Signal peptide} label using a Conditional Random Field.

**Figure 1:**
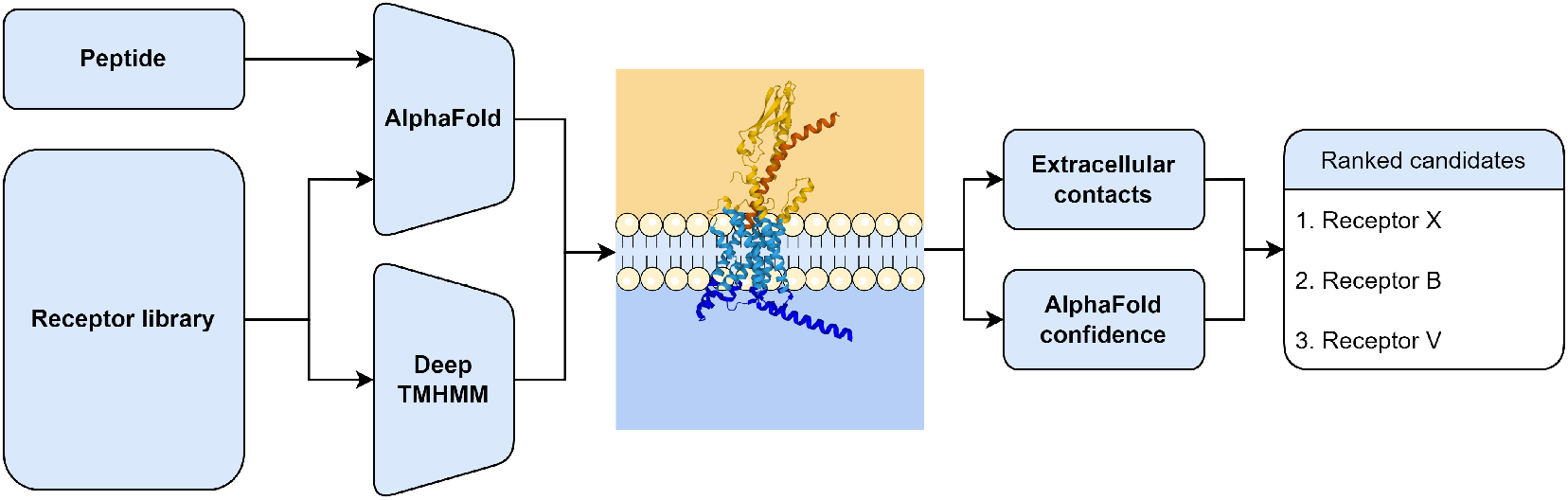
Peptide receptor ranking using AlphaFold-Multimer and DeepTMHMM. AlphaFold predicts a complex for each receptor with the peptide, DeepTMHMM the transmembrane topology of the receptor. Structure prediction confidence metrics and peptide-receptor contacts on the extracellular side are extracted and used to rank the candidate receptors.

We extract multiple prediction quality metrics from the predicted complex structure: median predicted aligned error at the interface (iPAE), median pLDDT of the peptide residues at the interface (ipLDDT) and the predicted interface TM score (ipTM). We also evaluate pDockQ [18], which was proposed for ranking protein-protein complexes, as a prediction quality score. For each metric, the receptors are ranked according to the score of their predicted complex. Using the topology, we extract the number of receptor residues labeled as extracellular that are in contact with the peptide. If this number is 0, we downrank the receptor as the peptide is predicted to bind at an intracellular interface or at sections that are buried in the cell membrane. These binding modes are biologically implausible, as they are not accessible to a peptide *in vivo*, but might still be predicted with high confidence by AlphaFold which has no explicit access to this prior information. For all interface calculations, we consider residues in contact if their distance is below 0.35 nm.

AlphaFold predicts five structures for each input, sampled from five different model checkpoints. For true binders, we expect all five predictions to have a similar predicted confidence. To penalize receptors that have a high variation, we aggregate the five predictions by taking the median and subtracting the median absolute deviation (MAD) (Equation 1). In the case of iPAE, where lower means better, we add the MAD. This corresponds to the 25^th^ percentile of the confidence distribution, thus favoring receptors with a narrow distribution.

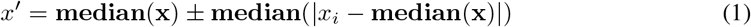

For each peptide-receptor pair, we report the percentile rank of the known receptor in the complete receptor library. Note that we do not necessarily expect the receptor to have the top rank - many peptides are known to have more than one endogenous binding partner [6]. Moreover, the exact ordering of receptors by AlphaFold confidence is unlikely to correlate to binding strength [19]. However, with respect to a full receptor library, any true receptor can still be expected to rank high.

## 3 Data

### 3.1 Benchmark peptide - receptor pairs

For benchmarking endogenous receptor identification performance, we gather structurally confirmed human peptide-receptor pairs that were released after the training data cutoff date of AlphaFold-Multimer using GPCRdb [7], yielding a total of 11 pairs (Table A1). To mimic the actual use case of only having sequence information available, we ignore any sequence modifications reported in the PDB and use the canonical UniProt [20] sequence for both the receptors and the peptides.

### 3.2 Membrane-bound receptors

We establish a human receptor library using UniProt annotations. We extract all proteins that are annotated with the keyword “Receptor [KW-0675]” and have a membrane subcellular location (“Cell membrane [SL-0039]”, “Cell surface [SL-0310]”) or the keyword “Transmembrane [KW-0812]”. From this set, we exclude all proteins that are additionally annotated with an intracellular membrane location (Mitochondrion, Nucleus, Acidocalcisome, Acrosome, COPI-coated vesicle, Endoplasmic reticulum-Golgi intermediate compartment membrane, Sarcoplasmic reticulum). We furthermore exclude single-pass transmembrane receptors, as they typically form multimers *in vivo* [21] and are often folded spuriously by AlphaFold if no further processing is applied [22]. Due to limited computational resources, we limit our study to proteins annotated in Swiss-Prot with a maximum length of 2000, yielding a total of 1112 receptor proteins.

### 3.3 Processing

We use the default AlphaFold-Multimer 2.2.0 pipeline with reduced databases to generate multiple sequence alignments (MSAs) for each peptide and each receptor sequence. Due to the combinatorial nature of the problem (a complex structure is predicted for each peptide with each receptor), precomputing the MSAs and assembling AlphaFold-Multimer inputs by combining the two MSAs before prediction results in a significant speedup, as MSA generation alone can take multiple hours. We do not use template information and omit the relaxation step.

## 4 Results

Out of all investigated confidence metrics, the predicted ipTM score is best suited for receptor ranking, followed by the iPAE and ipLDDT (Table 1). Overall, we reach a mean percentile rank of 0.69%, thereby ranking the known receptors higher than 99.31% of the library. In our library of 1112 receptors, this corresponds to the known receptor being contained within the top 8 candidates on average. We find that pDockQ, which was developed and validated on protein-protein complexes only, performs poorly for peptide-receptor binding ranking with a mean percentile rank of 11.51%, indicating that calibration on protein-protein complexes does not generalize to peptide-receptor interfaces. Using DeepTMHMM for downranking complexes with biologically implausible binding results in some improvement on average, with the effect varying greatly between different peptides. If the known pair is already scoring very high, DeepTMHMM has little to no effect. However, we observe that for some peptides, as exemplified by the case of Gastrin-17, many high-confidence implausible complexes are predicted. In such cases, DeepTMHMM application results in a strong improvement of the rank of the known receptor. In total, DeepTMHMM improved the rank of 7/12 peptide-receptor pairs.

**Table 1:** Percentile rank of true pairs using different ranking metrics. Lower means better.

| Peptide | Receptor | With DeepTMHMM [%] |  |  |  | Without DeepTMHMM [%] |  |  |  |
| --- | --- | --- | --- | --- | --- | --- | --- | --- | --- |
|  |  | ipTM | iPAE | ipLDDT | pDockQ | ipTM | iPAE | ipLDDT | pDockQ |
| Cholecystokinin-8 | CCKAR | **0.09** | 0.74 | 2.11 | 6.03 | **0.09** | 0.91 | 2.40 | 6.03 |
| Galanin | GALR1 | **0.09** | 0.10 | 0.10 | 12.86 | **0.09** | 0.10 | 0.10 | 29.77 |
| Galanin | GALR2 | **0.27** | 0.48 | 0.58 | 17.00 | 0.36 | 0.50 | 0.60 | 48.74 |
| Gastrin-17 | GASR | **1.08** | 2.82 | 7.87 | 9.71 | 16.73 | 28.21 | 49.30 | 46.40 |
| Ghrelin-27 | GHSR | **0.72** | 2.77 | 4.48 | 15.92 | 1.53 | 10.62 | 12.72 | 44.78 |
| Neuropeptide Y | NPY1R | **0.36** | 0.49 | 0.88 | 2.88 | **0.36** | 0.51 | 1.03 | 9.89 |
| Oxytocin | OXYR | **1.62** | 6.62 | 11.28 | 13.40 | 1.71 | 8.01 | 26.94 | 32.37 |
| Secretin | SECR | **0.36** | 1.29 | 0.50 | 9.44 | 0.45 | 7.24 | 0.64 | 43.79 |
| Somatoliberin | Q9HB45 | 0.63 | **0.39** | 0.58 | 9.53 | 0.63 | 0.42 | 0.84 | 29.59 |
| Somatostatin-14 | SSR2 | **0.09** | 0.10 | 0.69 | 15.29 | 0.36 | 0.32 | 1.28 | 40.92 |
| Substance P | NK1R | **2.25** | 3.12 | 8.75 | 14.57 | 2.43 | 4.79 | 16.23 | 30.58 |
| Mean |  | **0.69** | 1.72 | 3.44 | 11.51 | 2.25 | 5.60 | 10.19 | 32.99 |

Overall, although all known complexes were predicted correctly as per their DockQ [23] scores (Figure A1), we find that the distribution of prediction confidence scores varies greatly between different peptides (Figure 2). This prevents us from evaluating the presented approach as a classifier, as this would be done by pooling predictions at a given threshold over all the peptides and reporting a performance metric, which assumes that the score distributions are comparable. This confirms that when using just AlphaFold confidences without further downstream modeling, a ranking approach seems to be best suited for the problem.

**Figure 2:**
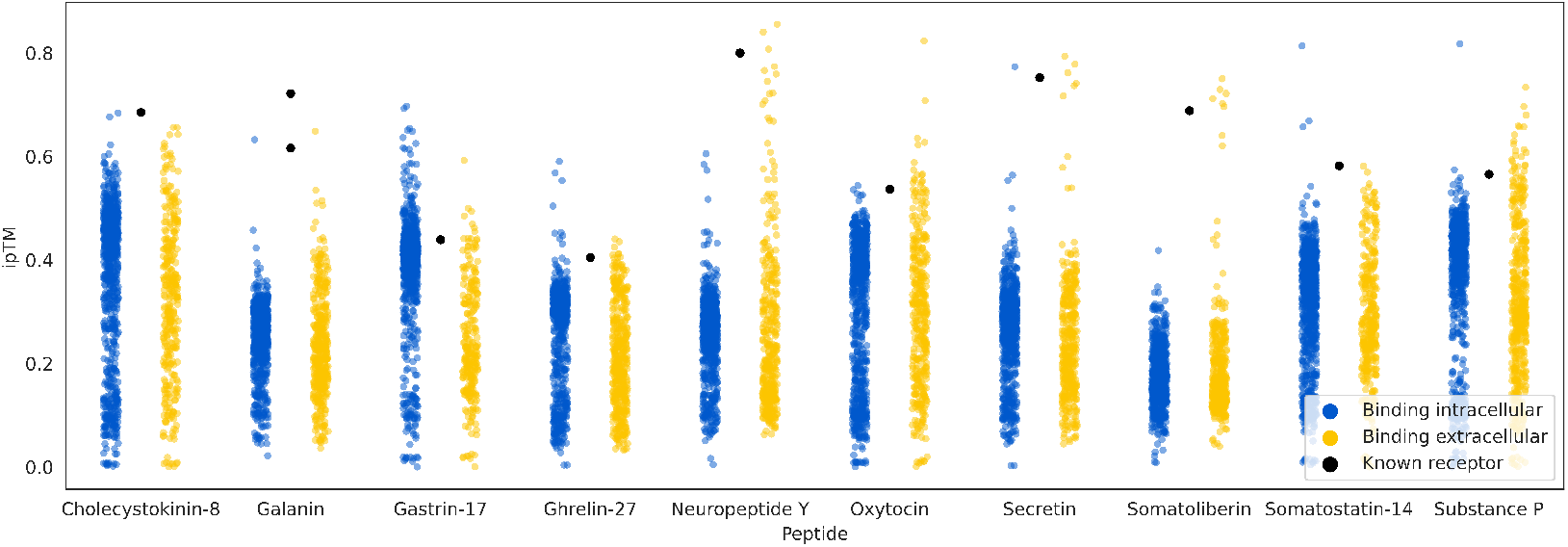
AlphaFold ipTM scores of all predicted peptide-receptor complexes of the test set. Each dot represents one complex. Intracellular-extracellular binding is computed using DeepTMHMM predicted topologies. All known receptors are predicted to bind on the extracellular side.

## 5 Discussion

Our results show that AlphaFold-Multimer can be applied successfully for prioritizing endogenous peptide receptors. Even though the proposed approach still requires a follow-up analysis to investigate the ranked list until the true receptor is found, it greatly reduces the complexity of such screening experiments. While conceivable, we refrained from developing a dedicated confidence-derived score such as pDockQ for peptide-receptor docking due to the very limited size of the dataset. Also, optimally, given a larger dataset, a classification model to distinguish true receptors could be trained on top of AlphaFold, as was recently done for peptide-MHC binding [24].

A key limitation of the presented approach is the computational demand of running AlphaFold on large libraries. Even though the MSA search becomes negligible when applying the combinatorial assembly strategy, the amount of required GPU hours is still a bottleneck that precludes using larger receptor libraries, such as including isoforms or multimeric receptors. The presented approach is easily adaptable to any other complex structure prediction method that outputs prediction confidences, so it will benefit from future performance improvements in the field [25, 26].

## Acknowledgments and Disclosure of Funding

We would like to thank Kristine Deibler for helpful discussions. We also thank Robin Andersson for generously providing GPU resources.

FT’s work was funded in part by the Novo Nordisk Foundation through the Center for Basic Machine Learning Research in Life Science (NNF20OC0062606). OW acknowledges support from the Pioneer Centre for AI, DNRF grant number P1.

## Code and Data availability

The receptor library, benchmark peptides and processing code are available at https://github.com/fteufel/alphafold-peptide-receptors

## A Appendix

**Table A1:**
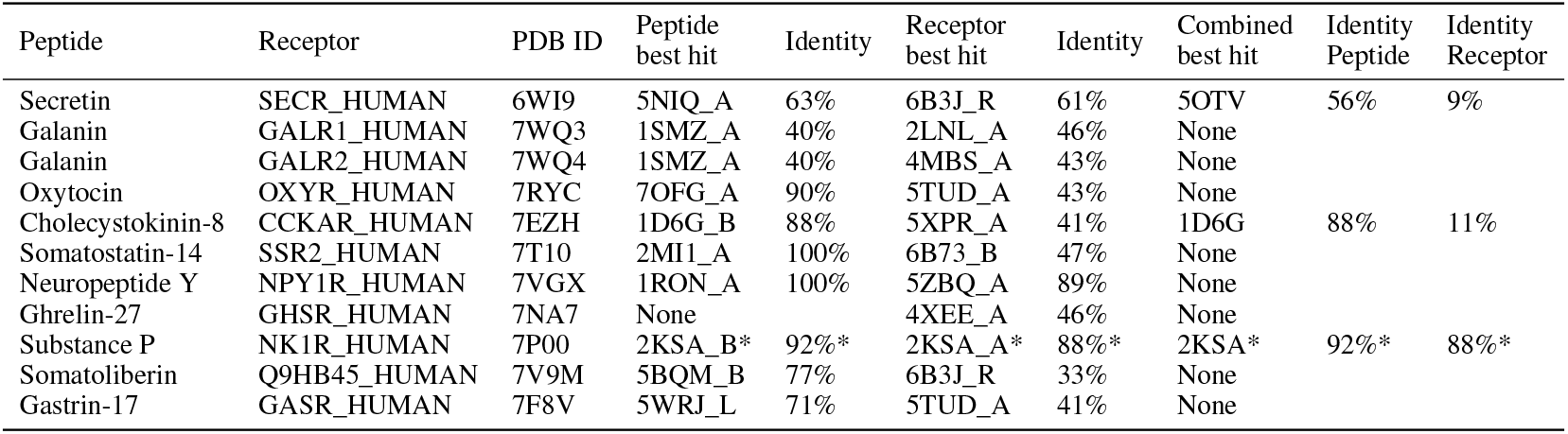
Peptide-Receptor complexes used in the benchmark experiment. To ensure that our benchmark complexes are not contained in the AlphaFold-Multimer training data as another PDB ID, we use BlastP to search the PDB for both the receptor and the peptide sequence and report their closest hits before the training cutoff date. The combined closest hit structure is determined by combining the ranks of each PDB ID in the peptide and receptor Blast hit lists. Sequence identities are computed with the length of the query sequence as denominator. For 7P00 there exist three earlier structures 2KS9, 2KSA, 2KSB representing the same sequences, but in a conformation that is different from 7P00 (DockQ scores 0.089, 0.083, 0.0103).

**Table A2:** Comparison of using the median and the MAD corrected median for pooling Alphafold prediction confidence metrics. All values are computed with DeepTMHMM filtering applied.

| Peptide | Receptor | Median | | | | Median ± MAD | | | |
| --- | --- | --- | --- | --- | --- | --- | --- | --- | --- |
|  |  | ipTM | iPAE | ipLDDT | pDockQ | ipTM | iPAE | ipLDDT | pDockQ |
| Cholecystokinin-8 | CCKAR | **0.09** | 0.29 | 3.24 | 6.47 | **0.09** | 0.74 | 2.11 | 6.03 |
| Galanin | GALR1 | **0.09** | **0.09** | 0.18 | 14.03 | **0.09** | 0.10 | 0.10 | 12.86 |
| Galanin | GALR2 | **0.27** | 0.46 | 0.74 | 18.17 | **0.27** | 0.48 | 0.58 | 17.00 |
| Gastrin-17 | GASR | 1.26 | 4.02 | 9.35 | 10.88 | **1.08** | 2.82 | 7.87 | 9.71 |
| Ghrelin-27 | GHSR | 1.17 | 2.80 | 5.41 | 17.00 | **0.72** | 2.77 | 4.48 | 15.92 |
| Neuropeptide Y | NPYR | 0.45 | 0.47 | 1.12 | 2.70 | **0.36** | 0.49 | 0.88 | 2.88 |
| Oxytocin | OXYR | 2.79 | 7.47 | 12.49 | 14.21 | **1.62** | 6.62 | 11.28 | 13.40 |
| Secretin | SECR | **0.36** | 0.66 | 0.66 | 9.44 | **0.36** | 1.29 | 0.50 | 9.44 |
| Somatoliberin | Q9HB45 | 0.63 | **0.19** | 1.03 | 14.3 | 0.63 | 0.39 | 0.58 | 9.53 |
| Somatostatin-14 | SSR2 | **0.09** | **0.09** | 0.95 | 16.10 | **0.09** | 0.10 | 0.69 | 15.29 |
| Substance P | NK1R | 2.97 | 4.49 | 10.32 | 15.38 | **2.25** | 3.12 | 8.75 | 14.57 |
| Mean |  | 0.92 | 1.91 | 4.13 | 12.61 | **0.69** | 1.71 | 3.44 | 11.51 |

**Table A3:** Overview of known peptide receptors that rank higher than the receptor included in the benchmark set. Receptors were not considered for the benchmark set if no experimental structure is available or the experimental structure was part of the AlphaFold-Multimer training data.

| Peptide | Receptor | Rank | Rank of benchmark set receptor |
| --- | --- | --- | --- |
| Somatoliberin | GHRHR_HUMAN | 1 | 7 |
| Neuropeptide Y | NPY2R_HUMAN | 1 | 4 |
| Neuropeptide Y | NPY5R_HUMAN | 2 | 4 |
| Galanin | GALR3_HUMAN | 2 | 1,3 |

**Figure A1:**
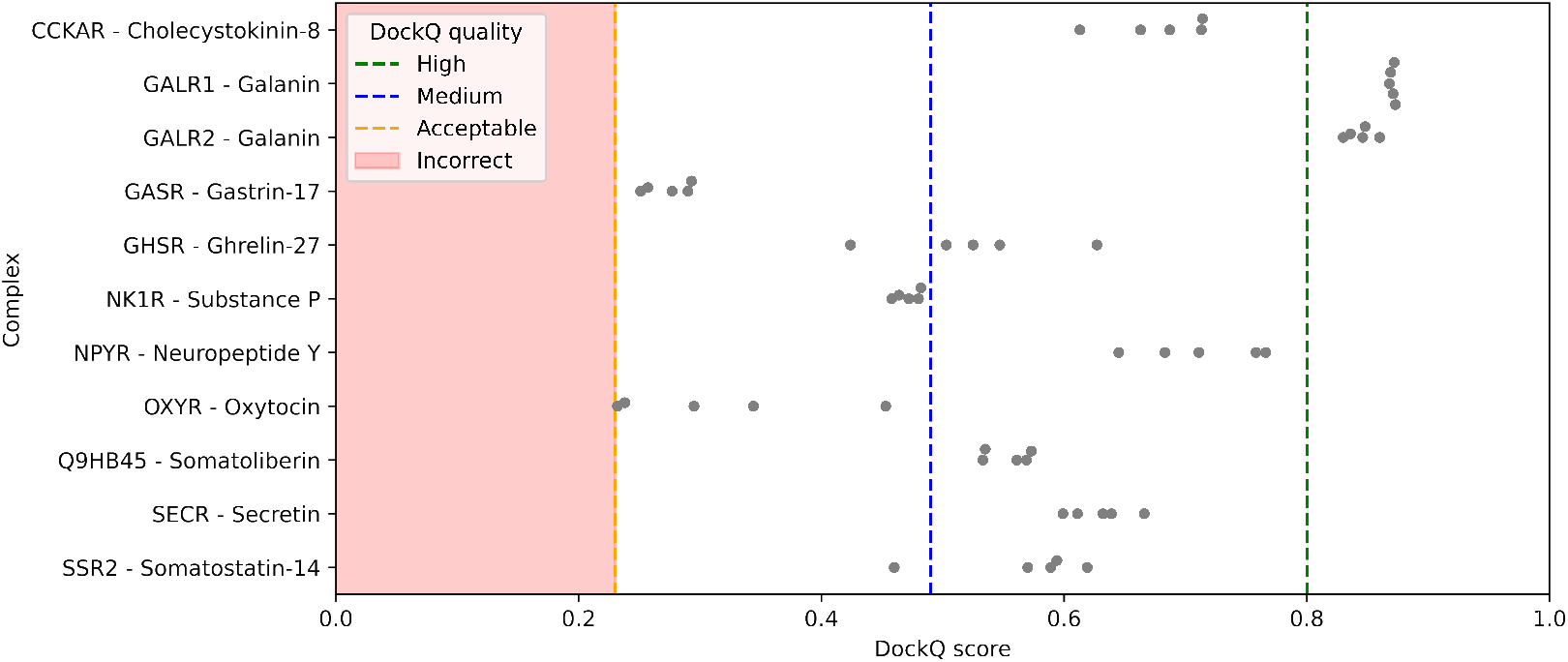
DockQ scores of the predicted AlphaFold-Multimer complexes compared to their experimental structures listed in Table A1.

## Notes

### Competing Interest Statement

The authors have declared no competing interest.

## References

[1] Paweł A. Kołodziejski, Ewa Pruszyńska-Oszmałek, Tatiana Wojciechowicz, Maciej Sassek, Natalia Leciejewska, Mariami Jasaszwili, Maria Billert, Emilian Małek, Dawid Szczepankiewicz, Magdalena Misiewicz-Mielnik, Iwona Hertig, Leszek Nogowski, Krzysztof W. Nowak, Mathias Z. Strowski, and Marek Skrzypski. The Role of Peptide Hormones Discovered in the 21st Century in the Regulation of Adipose Tissue Functions. Genes, 12(5):756, May 2021.

[2] Andrew F. Russo. Overview of neuropeptides: awakening the senses? Headache, 57(Suppl 2):37–46, May 2017.

[3] Markus Muttenthaler, Glenn F. King, David J. Adams, and Paul F. Alewood. Trends in peptide drug discovery. Nature Reviews Drug Discovery, 20(4):309–325, April 2021. Number: 4 Publisher: Nature Publishing Group.

[4] Anna Secher, Christian D. Kelstrup, Kilian W. Conde-Frieboes, Charles Pyke, Kirsten Raun, Birgitte S. Wulff, and Jesper V. Olsen. Analytic framework for peptidomics applied to large-scale neuropeptide identification. Nature Communications, 7(1):11436, May 2016. Number: 1 Publisher: Nature Publishing Group.

[5] Alan Saghatelian and Juan Pablo Couso. Discovery and characterization of smORF-encoded bioactive polypeptides. Nature Chemical Biology, 11(12):909–916, December 2015. Number: 12 Publisher: Nature Publishing Group.

[6] Simon R. Foster, Alexander S. Hauser, Line Vedel, Ryan T. Strachan, Xi-Ping Huang, Ariana C. Gavin, Sushrut D. Shah, Ajay P. Nayak, Linda M. Haugaard-Kedström, Raymond B. Penn, Bryan L. Roth, Hans Bräuner-Osborne, and David E. Gloriam. Discovery of Human Signaling Systems: Pairing Peptides to G Protein-Coupled Receptors. Cell, 179(4):895–908.e21, October 2019.

[7] Albert J Kooistra, Stefan Mordalski, Gáspár Pándy-Szekeres, Mauricio Esguerra, Alibek Mamyrbekov, Christian Munk, György M Keseru, and David E Gloriam. GPCRdb in 2021: integrating GPCR sequence, structure and function. Nucleic Acids Research, 49(D1):D335–D343, January 2021.

[8] Markus Sällman Almén, Karl JV Nordström, Robert Fredriksson, and Helgi B. Schiöth. Mapping the human membrane proteome: a majority of the human membrane proteins can be classified according to function and evolutionary origin. BMC Biology, 7(1):50, August 2009.

[9] Damaris Bausch-Fluck, Ulrich Goldmann, Sebastian Müller, Marc van Oostrum, Maik Müller, Olga T. Schubert, and Bernd Wollscheid. The in silico human surfaceome. Proceedings of the National Academy of Sciences, 115(46):E10988–E10997, November 2018. Publisher: Proceedings of the National Academy of Sciences.

[10] John Jumper, Richard Evans, Alexander Pritzel, Tim Green, Michael Figurnov, Olaf Ronneberger, Kathryn Tunyasuvunakool, Russ Bates, Augustin Žídek, Anna Potapenko, Alex Bridgland, Clemens Meyer, Simon A. A. Kohl, Andrew J. Ballard, Andrew Cowie, Bernardino Romera-Paredes, Stanislav Nikolov, Rishub Jain, Jonas Adler, Trevor Back, Stig Petersen, David Reiman, Ellen Clancy, Michal Zielinski, Martin Steinegger, Michalina Pacholska, Tamas Berghammer, Sebastian Bodenstein, David Silver, Oriol Vinyals, Andrew W. Senior, Koray Kavukcuoglu, Pushmeet Kohli, and Demis Hassabis. Highly accurate protein structure prediction with AlphaFold. Nature, 596(7873):583–589, August 2021. Number: 7873 Publisher: Nature Publishing Group.

[11] Richard Evans, Michael O’Neill, Alexander Pritzel, Natasha Antropova, Andrew Senior, Tim Green, Augustin Žídek, Russ Bates, Sam Blackwell, Jason Yim, Olaf Ronneberger, Sebastian Bodenstein, Michal Zielinski, Alex Bridgland, Anna Potapenko, Andrew Cowie, Kathryn Tunyasuvunakool, Rishub Jain, Ellen Clancy, Pushmeet Kohli, John Jumper, and Demis Hassabis. Protein complex prediction with AlphaFold-Multimer. bioRxiv, page 2021.10.04.463034, March 2022. Publisher: Cold Spring Harbor Laboratory Section: New Results.

[12] Isak Johansson-Åkhe and Björn Wallner. Improving Peptide-Protein Docking with AlphaFold-Multimer using Forced Sampling. bioRxiv, page 2021.11.16.468810, May 2022. Publisher: Cold Spring Harbor Laboratory Section: New Results.

[13] Tomer Tsaban, Julia K. Varga, Orly Avraham, Ziv Ben-Aharon, Alisa Khramushin, and Ora Schueler-Furman. Harnessing protein folding neural networks for peptide–protein docking. Nature Communications, 13(1):176, January 2022. Number: 1 Publisher: Nature Publishing Group.

[14] Henrik Nielsen, Konstantinos D. Tsirigos, Søren Brunak, and Gunnar von Heijne. A Brief History of Protein Sorting Prediction. The Protein Journal, 38(3):200–216, June 2019.

[15] Tamás Hegedus, Markus Geisler, Gergely László Lukács, and Bianka Farkas. Ins and outs of AlphaFold2 transmembrane protein structure predictions. Cellular and Molecular Life Sciences, 79(1):73, January 2022.

[16] Jeppe Hallgren, Konstantinos D. Tsirigos, Mads Damgaard Pedersen, José Juan Almagro Armenteros, Paolo Marcatili, Henrik Nielsen, Anders Krogh, and Ole Winther. DeepTMHMM predicts alpha and beta transmembrane proteins using deep neural networks. Technical report, bioRxiv, April 2022. Section: New Results Type: article.

[17] Alexander Rives, Joshua Meier, Tom Sercu, Siddharth Goyal, Zeming Lin, Jason Liu, Demi Guo, Myle Ott, C. Lawrence Zitnick, Jerry Ma, and Rob Fergus. Biological structure and function emerge from scaling unsupervised learning to 250 million protein sequences. Proceedings of the National Academy of Sciences, 118(15):e2016239118, April 2021. Publisher: Proceedings of the National Academy of Sciences.

[18] Patrick Bryant, Gabriele Pozzati, and Arne Elofsson. Improved prediction of protein-protein interactions using AlphaFold2. Nature Communications, 13(1):1265, March 2022. Number: 1 Publisher: Nature Publishing Group.

[19] Patrick Bryant and Arne Elofsson. EvoBind: in silico directed evolution of peptide binders with AlphaFold. bioRxiv, page 2022.07.23.501214, July 2022. Publisher: Cold Spring Harbor Laboratory Section: New Results.

[20] The UniProt Consortium. UniProt: the universal protein knowledgebase in 2021. Nucleic Acids Research, 49(D1):D480–D489, January 2021.

[21] Katrine Bugge, Kresten Lindorff-Larsen, and Birthe B. Kragelund. Understanding single-pass transmembrane receptor signaling from a structural viewpoint—what are we missing? The FEBS Journal, 283(24):4424–4451, 2016. _eprint: https://onlinelibrary.wiley.com/doi/pdf/10.1111/febs.13793.

[22] Andrei L. Lomize, Kevin A. Schnitzer, Spencer C. Todd, Stanislav Cherepanov, Carlos Outeiral, Charlotte M. Deane, and Irina D. Pogozheva. Membranome 3.0: Database of single-pass membrane proteins with AlphaFold models. Protein Science, 31(5):e4318, 2022. _eprint: https://onlinelibrary.wiley.com/doi/pdf/10.1002/pro.4318.

[23] Sankar Basu and Björn Wallner. DockQ: A Quality Measure for Protein-Protein Docking Models. PLOS ONE, 11(8):e0161879, August 2016. Publisher: Public Library of Science.

[24] Amir Motmaen, Justas Dauparas, Minkyung Baek, Mohamad H. Abedi, David Baker, and Philip Bradley. Peptide binding specificity prediction using fine-tuned protein structure prediction networks, July 2022. Pages: 2022.07.12.499365 Section: New Results.

[25] Ruidong Wu, Fan Ding, Rui Wang, Rui Shen, Xiwen Zhang, Shitong Luo, Chenpeng Su, Zuofan Wu, Qi Xie, Bonnie Berger, Jianzhu Ma, and Jian Peng. High-resolution de novo structure prediction from primary sequence. bioRxiv, page 2022.07.21.500999, July 2022. Publisher: Cold Spring Harbor Laboratory Section: New Results.

[26] Zeming Lin, Halil Akin, Roshan Rao, Brian Hie, Zhongkai Zhu, Wenting Lu, Allan dos Santos Costa, Maryam Fazel-Zarandi, Tom Sercu, Sal Candido, and Alexander Rives. Language models of protein sequences at the scale of evolution enable accurate structure prediction. bioRxiv, page 2022.07.20.500902, July 2022. Publisher: Cold Spring Harbor Laboratory Section: New Results.

