## Supplementary figures and images for "Identifying endogenous peptide receptors by combining structure and transmembrane topology prediction"

### Supplementary Figure 1

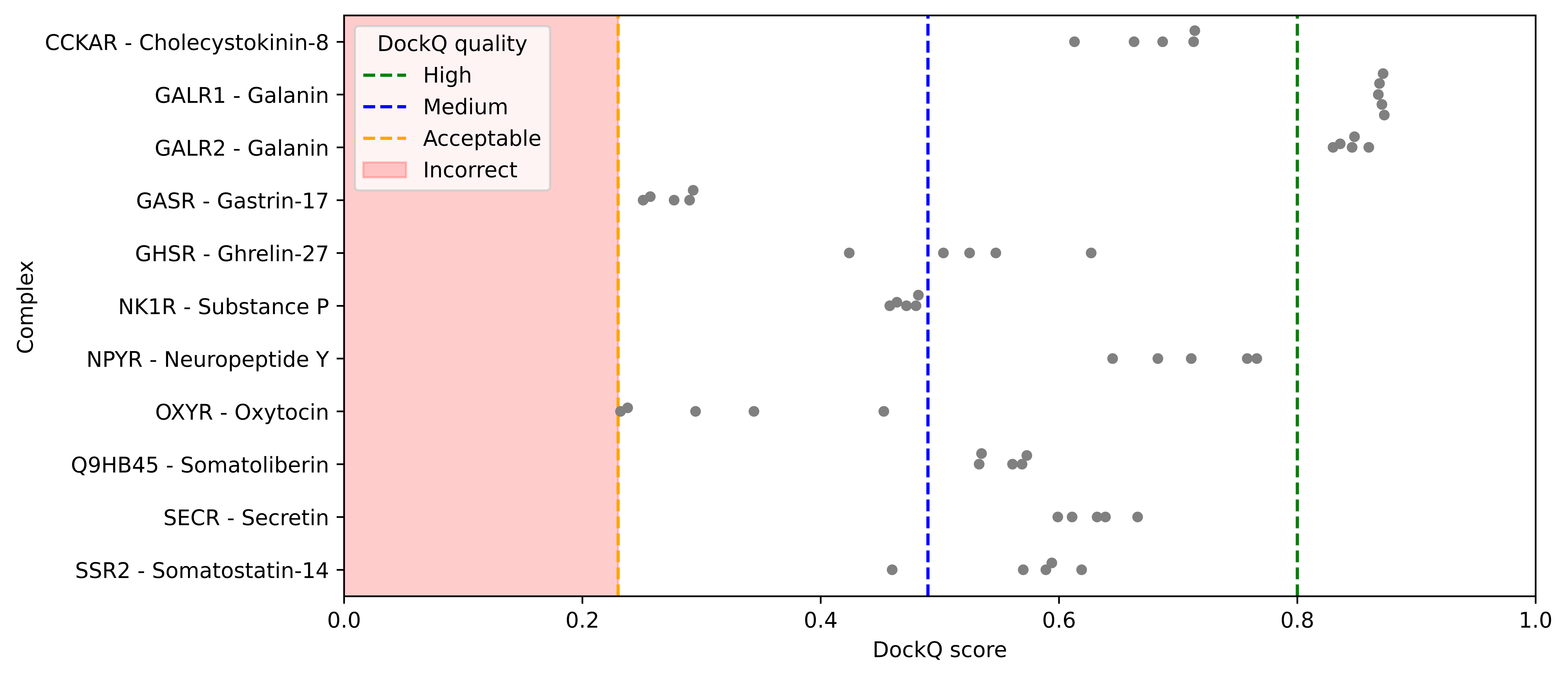
